# Locally secreted BiTEs complement CAR T cells by enhancing solid tumor killing

**DOI:** 10.1101/2021.06.08.447602

**Authors:** Yibo Yin, Jesse L. Rodriguez, Nannan Li, Radhika Thokala, MacLean P. Nasrallah, Li Hu, Logan Zhang, Jiasi Vicky Zhang, Devneet Kainth, Leila Haddad, Emily X. Johns, Yu Long, Hongsheng Liang, Jiping Qi, Xiangtong Zhang, Zev A. Binder, Zhiguo Lin, Donald M. O’Rourke

## Abstract

Bispecific T-cell engagers (BiTEs) are bispecific antibodies that redirect T cells to target antigen-expressing tumors. BiTEs can be secreted by T cells through genetic engineering and perform anti-tumor activity. We hypothesized that BiTE-secreting T cells could be a valuable T cell-directed therapy in solid tumors, with distinct properties in mono- or multi-valent strategies incorporating chimeric antigen receptor (CAR) T cells. Glioblastomas represent a good model for solid tumor heterogeneity and represent a significant therapeutic challenge. We detected expression of tumor-associated epidermal growth factor receptor (EGFR), EGFR variant III (EGFRvIII), and interleukin-13 receptor alpha 2 (IL13Rα2) on glioma tissues and glioma cancer stem cells. These antigens formed the basis of a multivalent approach, using a conformation-specific tumor-related EGFR targeting antibody (806) and Hu08, an IL13Rα2-targeting antibody, as the scFvs to generate new BiTE molecules. Compared with 806CAR T cells and Hu08CAR T cells, BiTE T cells demonstrated prominent activation, cytokine production, and cytotoxicity in response to target-positive gliomas. Superior response activity was also demonstrated in BiTE secreting bivalent targeting T cells compared with bivalent targeting CAR T cells, which significantly delayed tumor growth in a glioma mouse model. In summary, BiTEs secreted by mono- or multi- valent targeting T cells have potent anti-tumor activity *in vitro* and *in vivo* with significant sensitivity and specificity, demonstrating a promising strategy in solid tumor therapy.

## INTRODUCTION

T cells can be redirected to target tumors by being genetically modified to express chimeric antigen receptors (CARs) (*1*). To date, mixed success has been obtained in liquid and solid tumors using this approach. Sustained remission was achieved in leukemia and lymphoma patients treated with CD19 targeting CAR T cells (*2–4*). 85% of patients with relapsed or refractory multiple myeloma responded to B-cell maturation antigen targeting (*5*). In the realm of solid tumors, clinical application of CAR T cells has had limited success. A patient with recurrent multifocal glioblastoma (GBM) also had a 7.5-month clinical response after the treatment of interleukin-13 receptor alpha 2 (IL13Rα2) targeting CAR T cells (*6*). This patient’s tumor recurred with low levels of the target antigen. This underscores the potential use of CAR T cells against solid tumors, in addition to bringing to light the limitations of monotherapy approaches against dynamic tumors such as GBM.

Another approach to redirect T cells against tumors is bispecific T cell engagers (BiTEs). BiTEs combine antigen specificity with the ability to induce cytotoxicity via bystander T cells by linking two single chain variable fragments (scFvs), one recognizing a tumor antigen and the other recognizing the CD3 molecule on bystander T cells (*7*). Patients with leukemia or lymphoma treated with a CD19-targeting BiTE (blinatumomab) achieved sustained remission (*8–11*). However, repeated infusions were needed, as the half-life of BiTEs in circulation was approximately 2-3 hours (*7*). In a previous study, T cells transduced to secrete an EphA2-targeting BiTE were shown to successfully activate bystander T cells and inhibit tumor growth in glioma and lung cancer murine models (*12*). In another report directly comparing the anti-tumor activity of mRNA electroporated blinatumomab-secreting T cells and CD19 CAR T cells, the BiTE T cells demonstrated superior tumor suppression (*13*).

Despite the prominent effect achieved by CAR T cells and BiTEs in the treatment of leukemia and lymphoma, a significant number of treated patients have had limited antitumor activity or tumor relapse (*1, 7, 14*). Antigen loss or downregulation is a common mechanism of treatment resistance, which could be mitigated by combinatorial targeting of multiple antigens (*15–17*). Antigen loss was also observed in solid tumors after CAR T cell therapy. Significant antigen loss was seen in recurrent GBM patients after the treatment of IL13Rα2-targeting CAR T cells and epidermal growth factor receptor (EGFR) variant III (EGFRvIII)-targeting CAR T cells (*6, 18*). Preclinical work on monovalent BiTEs in GBM have included EGFRvIII-targeting (*19*) and IL13Rα2-targeting (*20*) constructs. To address antigen heterogeneity in solid tumors, multivalent CAR T cells have been generated and studied in pre-clinical models, allowing for expansion of targetable tumors and amplification of treatment effects (*21–23*). EGFR-targeting BiTEs secreted from EGFRvIII-targeting CAR T cells have also been developed preclinically. They demonstrated the feasibility of delivering EGFR-targeting BiTEs via EGFRvIII-targeting CAR T cells (*24*), which suggests BiTEs can be also utilized in generating multivalent T cells.

GBMs are the most common primary malignant brain tumor (*25, 26*). With standard-of-care treatment, including surgical resection, radiotherapy, chemotherapy, and tumor-treating fields, the median survival is only 12-17 months (*26, 27*). Single-cell RNA sequencing revealed tremendous intra-tumoral heterogeneity within GBM (*28*). In our phase I trial of EGFRvIII-targeting CAR T cells, we found EGFRvIII expression was downregulated post-CAR T infusion, but not amplified wild-type EGFR (*18*). Independently, IL13Rα2 was reported to cooperate with EGFRvIII in promoting GBM tumorigenicity (*29*). We hypothesized the simultaneous targeting of these two antigens may result in a meaningful clinical impact for patients with GBM. We have previously reported on the anti-tumor activity of an IL13Rα2-specific CAR (*30*). Our group and others have also demonstrated the anti-tumor activity of a pan-EGFR alteration-specific CAR based on the 806 monoclonal antibody (mAb) (*31, 32*). The 806 mAb recognizes a cryptic epitope on EGFR that is exposed in mutated or amplified conformations found on tumors (*33, 34*). 806 does not exhibit high affinity for physiologic EGFR, due to differences in mannose modification when expressed at normal levels (*35*). In this study, we generated 806BiTE and Hu08BiTE secreting T cells, and compared activity to 806CAR T cells and Hu08CAR T cells in responding to GBM cells. We demonstrate the anti-tumor activity, sensitivity, and specificity in mono- and multi-valent targeting constructs that have application to GBM and other solid tumors.

## RESULTS

### EGFRvIII co-existed with EGFR and IL13Rα2 in GBM

We performed flow cytometry on freshly resected glioma tissue to demonstrate the expression of EGFR and EGFRvIII in glioma cells (Figure 1A). The expression of both targets was confirmed, with cases demonstrating significantly more EGFR positivity than EGFRvIII (p=0.0172). Staining results of four cases with the highest EGFRvIII expression illustrated that a considerable percentage (10.6%, 22.7%, 14.9% and 34.1%) of EGFR positive cells were EGFRvIII negative. This suggested targeting EGFR would cover a greater percentage of tumor cells than targeting EGFRvIII. Case 8739, being the exception, had more than half of its tumor cells negative for both EGFR and EGFRvIII, highlighting the need for additional targets beyond EGFR variants.

**Fig. 1.**
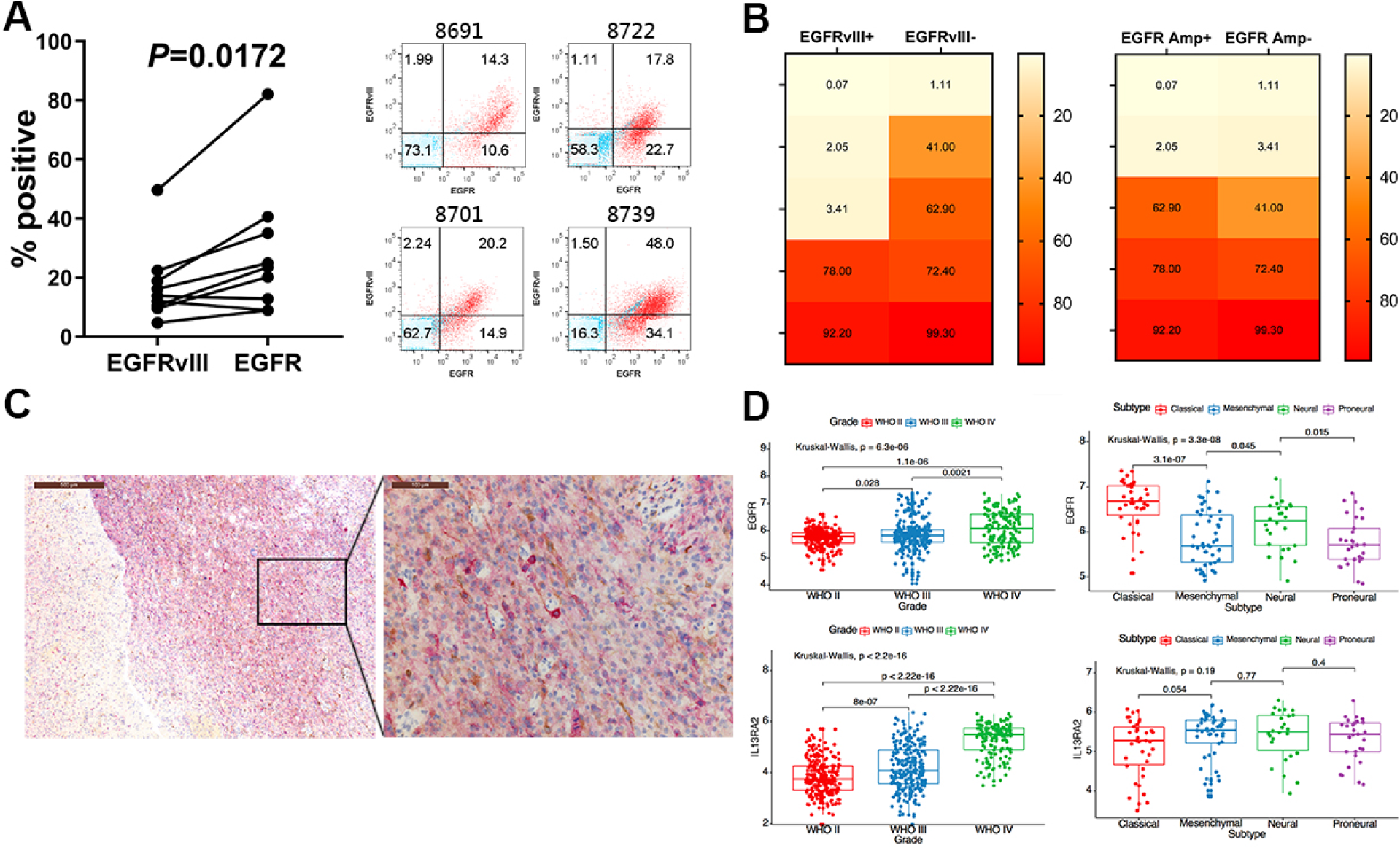
EGFR, EGFRvIII, and IL13Rα2 co-express in GBM. (A) Flow cytometry analyses of EGFRvIII and EGFR expression in resected glioma tissues (left). Statistically significant differences were determined using two-tailed paired t test. Flow panels indicated EGFR expression along x-axis and EGFRvIII expression along y-axis of top four EGFRvIII expression cases (right). (B) Heat maps showing percentage of IL13Rα2 positive cells in EGFRvIII positive or negative (left) and amplified or unamplified EGFR (right) GSC lines. (C) Immunohistochemical stains of IL13Rα2 (brown) and EGFRvIII (red) in resected glioma tissues. (D) EGFR (upper) and IL13Rα2 (lower) expression segregated by WHO grades (left) and GBM molecular subtypes (right) based on the RNA-seq data from the Cancer Genome Atlas (TCGA) database. Statistically significant differences were calculated by Kruskal-Wallis Test with p<0.05 being considered statistically significant.

To broaden the range of targetable tumor cells, we also determined IL13Rα2 expression in ten glioma stem cell (GSC) lines from cases with varying levels EGFR and EGFRvIII expression as determined by next generation sequencing (Figure 1B) (*36*). The expression of IL13Rα2 was also found to be heterogeneous throughout the tumor. Six out of ten cases had more than 40% detectable expression of IL13Rα2. In these six cases, two were positive for EGFRvIII expression and three were positive for EGFR amplification. In the four IL13Rα2-negative cases, only one was negative for both EGFR amplification and EGFRvIII expression. We also detected IL13Rα2 co-expressed with EGFRvIII in glioma tissues by immunohistochemistry (IHC) (Figure 1C). Significant heterogeneity of both targets was observed in the same tissue slice. These results suggested that the addition of IL13Rα2 as a target would expand the application of EGFR and EGFRvIII targeting strategies.

In order to analyze target expression in a larger patient population, we downloaded RNA-seq data of gliomas from the Genomic Data Commons data portal on The Cancer Genome Atlas (TCGA). Glioma cases were grouped by both WHO grade and genomic subtype (*37*) (Figure 1D). The expression of EGFR and IL13Rα2 was significantly higher in high grade gliomas than low grade gliomas. EGFR was predominately expressed in the Classical and Neural subtypes, with lower expression levels in the Mesenchymal and Proneural subtypes. There was no statistical difference in IL13Rα2 expression between glioma subtypes. These results also demonstrated the existence of EGFR and IL13Rα2 in glioma tissue and the significance of targeting both antigens with T cell engaging therapies.

### BiTEs secreted from T cells bind targets with a high degree of specificity

Given the heterogeneous antigen expression found in gliomas, we next set out to determine if BiTE-secreting T cells could be used as an antigen-specific targeting strategy. We generated an EGFR-targeting BiTE and an IL13Rα2-targeting BiTE with the 806 and Hu08 scFvs, respectively. Both 806CAR and Hu08CAR constructs were also included to compare the anti-tumor activity of the corresponding BiTE constructs. mCherry was co-expressed with each construct by a self-cleaving sequence (T2A) to demonstrate transduction efficacy (Figure 2A).

**Fig. 2.**
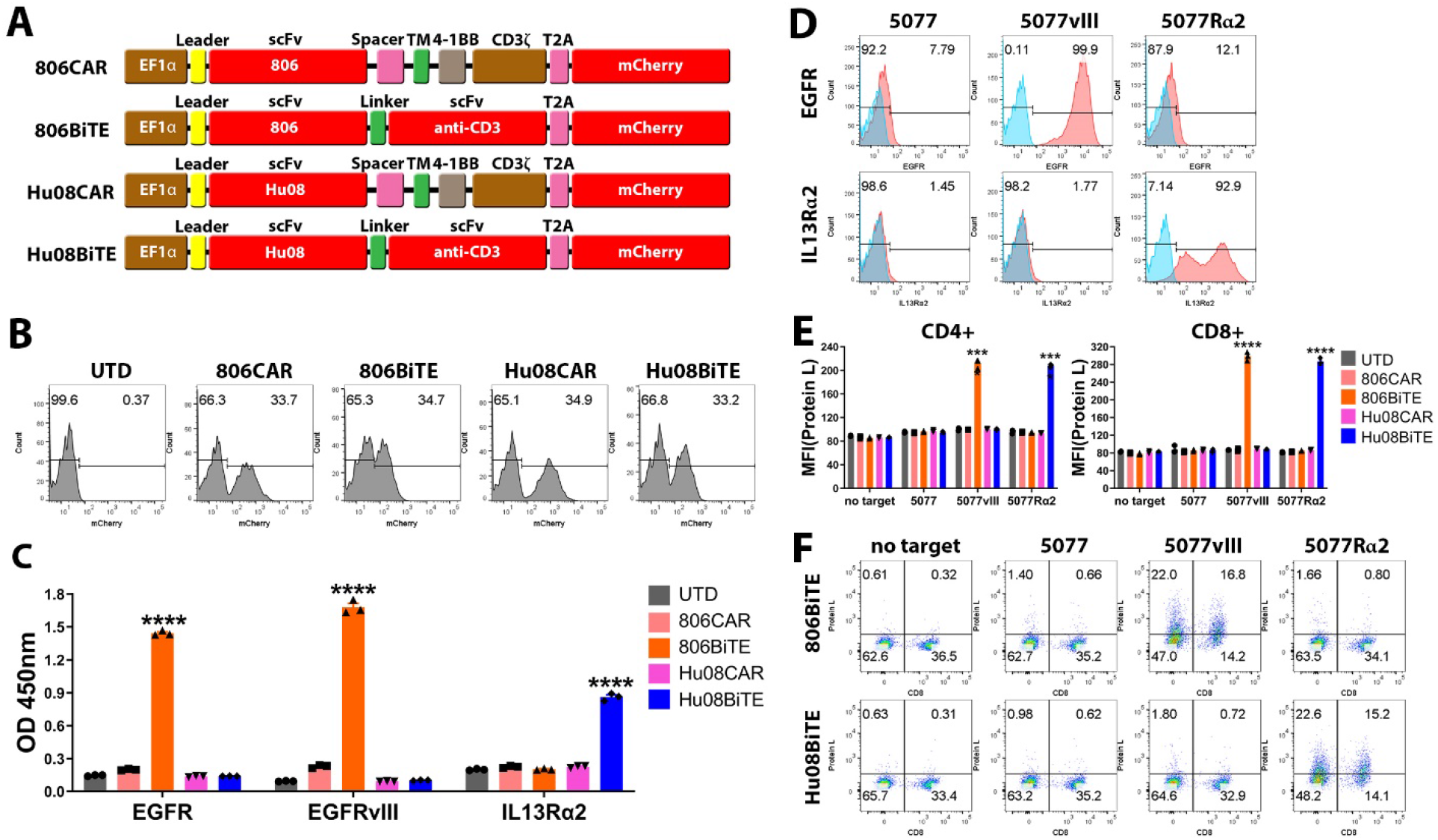
BiTEs secreted from T cells specifically bind to target antigens and T cells. (A) Schematic vector maps of 806CAR/806BiTE/Hu08CAR/Hu08BiTE constructs. (B) Flow cytometric detection of T cell transduction by mCherry expression. (C) Conditioned media of T cells in each group tested by ELISA to evaluate the binding ability of secreted BiTEs to recombinant EGFR, EGFRvIII and IL13Rα2 proteins. (D) The GSC line 5077 was lentivirally-transduced to overexpress EGFRvIII or IL13Rα2 and assessed via flow cytometry. (E) Conditioned media of CAR/BiTE T cells was collected and co-cultured with UTD T cells and target cells. BiTE binding on T cells was detected by biotinylated protein L with secondary streptavidin coupled FITC after 16hrs co-culture. The median fluorescence intensity (MFI) was quantified on CD4 and CD8 positive T cells. (F) Flow based results of representative samples in (E). CD8 was stained to distinguish the CD4-positive and CD8-positive subgroups of T cells along the x axis. Statistically significant differences were calculated by one-way Analysis of Variance (ANOVA) with post hoc Tukey test. ****p<0.0001. Data are presented as means ± SEM.

After transduction with each construct, mCherry expression was detected on T cells by flow cytometry, indicating similar transduction efficacy (Figure 2B). Conditioned media from each group was collected and used in a standard direct ELISA to confirm the secretion and binding of 806BiTE and Hu08BiTE from T cells to recombinant EGFR, EGFRvIII, and IL13Rα2 (Figure 2C). 806BiTE secreted from T cells significantly detected plate-bound EGFR and EGFRvIII (p<0.0001). The OD value in the EGFRvIII binding group was higher than EGFR binding group, suggesting a high affinity of the 806BiTE for EGFRvIII (p=0.0006). Hu08BiTE secreted from T cells also significantly bound to IL13Rα2 (p<0.0001) when compared with the other T-cell groups.

To further demonstrate the characteristics of BiTE T cells, we selected GSC line 5077 (*38*) which expressed low levels of endogenous EGFR (7.79%) but not IL13Rα2 or EGFRvIII. We engineered 5077 cells to overexpress EGFRvIII or IL13Rα2 as target cells (Figure 2D). Conditioned media from each BiTE T cell group was collected and used in co-culture with untransduced (UTD) T cells and target cells. BiTE binding to T cells was detected with protein L staining by flow cytometry, showing target-specific binding (Figure 2E). 806BiTE or Hu08BiTE binding on T cells was only detected when EGFRvIII or IL13Rα2 was overexpressed in 5077 cells, respectively (p<0.0001). No BiTE binding was detectable on T cells in the UTD T cell groups cultured without target cells or with the parental cell line. Flow based results of representative samples are illustrated (Figure 2F), indicating that a BiTE’s ability to bind to T cells was eliminated in the absence of target antigen.

### BiTEs and BiTE T cells respond to antigen positive glioma cells

To determine the ability of the secreted 806BiTEs and Hu08BiTEs to initiate T cell responses, T cell activation marker CD69 was assessed on UTD T cells by flow cytometry, when co-cultured with target cells in conditioned media from BiTE transduced T cells. Conditioned media from CAR T cells was used as a negative control (Figure 3A). CD69 expression on UTD T cells was significantly elevated in co-culture with 5077^EGFRvIII+^ cells with conditioned media from 806BiTE T cells and co-culture with 5077^IL13Rα2+^ cells with conditioned media from Hu08BiTE T cells (p<0.0001). CD69 expression elevation was noted in both CD4 and CD8 positive UTD T cells. Flow panels of representative samples indicated both 806BiTEs and Hu08BiTEs induced UTD T cell activation in the presence of antigen positive target cells (Figure 3B). Cytokine production (IFNγ, IL2, and TNFα) of UTD T cells was also detected when co-cultured with target cells in conditioned media (Figure S1), which was consistent with the result of CD69 expression on these UTD T cells. Only 5077^EGFRvIII+^ cells in co-culture with conditioned media from 806BiTE T cells and 5077^IL13Rα2+^ cells in co-culture with conditioned media from Hu08BiTE T cells induced significant cytokine production, in both CD4+ and CD8+ T cells subgroups (p<0.0001). No statistical differences were observed between any other groups. To determine the ability of 806BiTE and Hu08BiTE to mediate antigen-specific cytotoxicity, we performed bioluminescence cytotoxicity assays with UTD T cells co-cultured with different target cells. No cytotoxic activity was observed in co-cultures with 5077. Significant killing activity was detected in 5077^EGFRvIII+^ cells co-cultured with conditioned media from 806BiTE T cells and 5077^IL13Rα2+^ cells co-cultured with conditioned media from Hu08BiTE T cells. These results demonstrated that T cells transduced to secrete 806BiTEs or Hu08BiTEs can significantly and specifically activate UTD T cells in an antigen-specific manner (Figure 3C).

**Fig. 3.**
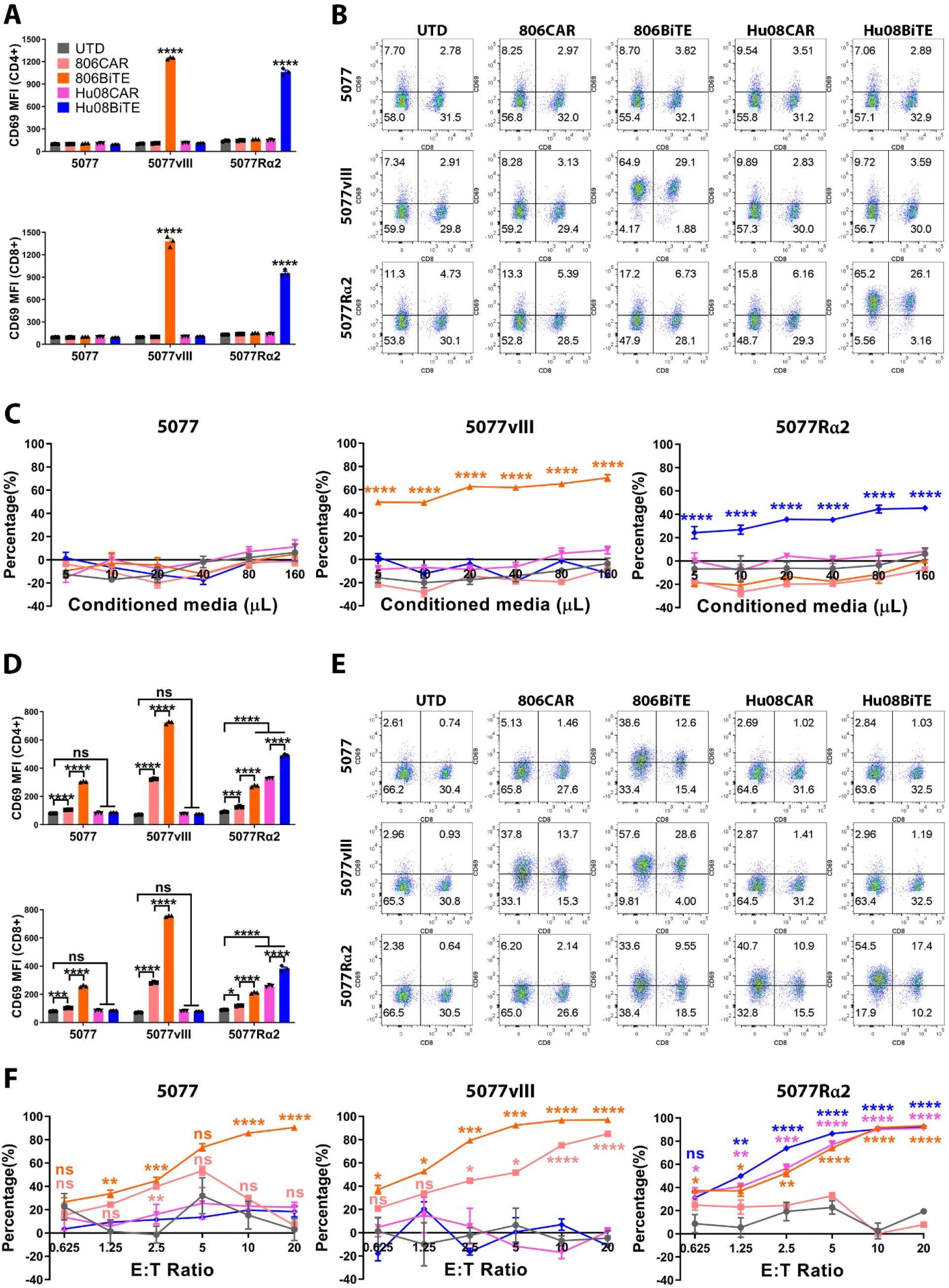
BiTEs and BiTE T cells respond to target positive glioma cells. (A) UTD T cell activation, as demonstrated by CD69 expression, in CD4+ (top) and CD8+ (bottom) T cells after 16hrs co-culture with conditioned media from CAR/BiTE T cells. (B) Flow based results of representative samples in (A). CD8 was stained to distinguish the CD4-positive and CD8-positive subgroups of T cells along the x axis. (C) Bioluminescence cytotoxicity assay of UTD T cells co-cultured with tumor cell lines and conditioned media of CAR/BiTE T cells. (D) T cell activation, as measured by CD69 expression, in CD4+ (top) and CD8+ (bottom) T cells after 16hrs co-culture of CAR/BiTE T cells with target cells. (E) Flow cytometry panels of bar graphs in (D). CD8 was stained to distinguish the CD4-positive and CD8-positive subgroups of T cells along the x axis. (F) Bioluminescence cytotoxicity assays of CAR/BiTE T cells co-cultured with tumor cell lines was analyzed at different effector/target (E:T) ratios (0.625:1, 1.25:1, 2.5:1, 5:1, 10:1, and 20:1) and compared with the UTD T cell group. Statistically significant differences were calculated by one-way ANOVA with post hoc Tukey test. ns, not significant; *p<0.05, **p<0.01, ***p<0.001, ****p<0.0001. Data are presented as means ± SEM.

Next, we considered the ability of 806BiTE T cells and Hu08BiTE T cells to respond to target-expressing tumor cells, 806BiTE T cells and Hu08BiTE T cells by co-culturing them with parental 5077, 5077^EGFRvIII+^, or 5077^IL13Rα2+^ cells. We included positive control 806CAR T cells and Hu08CAR T cells and negative control UTD T cells. Both 806BiTE T cells and Hu08BiTE T cells significantly activated by 5077^EGFRvIII+^ and 5077^IL13Rα2+^ cells, respectively, in both CD4+ and CD8+ T cells subgroups compared with UTD groups (p<0.0001). The stimulation level was even higher than 806CAR T cells and Hu08CAR T cells in co-culture with the same target cells (p<0.0001). Interestingly, CD69 expression on 806BiTE T cells and 806CAR T cells was also elevated when co-cultured with parental 5077 and 5077^IL13Rα2+^ cells in CD4+ and CD8+ T cells subgroups, which was likely due to the detection of the background EGFR expression in parental 5077 cells. The activation level of 806BiTE T cells was higher than that of 806CAR T cells in both CD4+ and CD8+ T cell subgroups (p<0.0001) (Figure 3D and 3E). To further determine the ability of 806BiTE and Hu08BiTE T cells to mediate antigen-specific cytotoxicity, bioluminescence cytotoxicity assays were performed by co-culturing 806 and Hu08 BiTE/CAR T cells with parental 5077, 5077^EGFRvIII+^, or 5077^IL13Rα2+^ cells (Figure 3F) in fresh media for 18hrs. Consistent with increased T cell activation of 806BiTE T cells co-cultured with 5077, significant cytotoxicity was also observed when 806BiTE secreting T cells were co-cultured with 5077 cells. This was not observed in UTD T cells co-cultured with 5077 in the presence of 806BiTE conditioned media, indicating that autocrine binding of 806BiTE to T cells results in the ability of these T cells to respond to low antigen expressing tumor cells. 806CAR T cells did not induce cytotoxicity when co-cultured with 5077 cells, nor did Hu08CAR T cells or Hu08BiTE T cells. In 5077^EGFRvIII+^ co-culture groups, high levels of cytotoxicity by both 806CAR and 806BiTE T cells were observed. 806BiTE T cells exhibited higher cytotoxicity than 806CAR T cells at 5:1 and 10:1 effector to target ratios (p<0.05). In 5077^IL13Rα2+^ co-culture groups, both Hu08CAR T cells and Hu08BiTE T cells demonstrated potent cytotoxicity. 806BiTE T cells also induced cytotoxicity, which was again consistent with the results of T cell activation and cytotoxicity assays when these cells were co-cultured with 5077 cells. This result was likely due to low levels of EGFR expressed on 5077 parental cells that were targeted by 806BiTE T cells.

### BiTE T cells respond to low target antigen expression on glioma cancer stem cells

To determine the sensitivity of 806BiTE to detect low levels of EGFR expression on 5077 cells, we generated 806BiTE T cells with high (H), medium (M), and low (L) levels of transduction efficacy. 806CAR T cells with comparable transduction efficacy were included as control (Figure 4A). Consistent with previous results, CD69 expression on T cells was upregulated in both 806BiTE and 806CAR T cell groups when co-cultured with 5077 cells (Figure 4B). The upregulated levels were concordant with transduction efficacy. CD69 expression was significantly higher in 806BiTE transduced T cells than 806CAR transduced T cells (p<0.0001 for both CD4+ and CD8+ subgroups at high and medium levels of transduction, p=0.0003 and p=0.3554 in CD4+ and CD8+ subgroups of low levels of transduction, respectively). Conditioned media of 806BiTE T cells and 806CAR T cells at different transduction levels was collected and added in UTD T cells co-cultured with 5077. No significant difference of cytotoxicity in any groups was detected as compared with conditioned media added from the UTD T cell group (Figure 4C, left panel). When 806BiTE T cells at different transduction levels were co-cultured with 5077 cells in fresh media, cytotoxicity was observed in every group in a dose-dependent manner (Figure 4C, right panel). No cytotoxicity was observed in 806CAR T cells co-cultured with 5077 at any transduction level. These results demonstrated a dose-dependent activation and killing activity of 806BiTE T cells co-cultured with 5077, expressing low levels of endogenous EGFR, which was not observed in 806CAR T cells or 806BiTE recruited UTD T cells.

**Figure 4.**
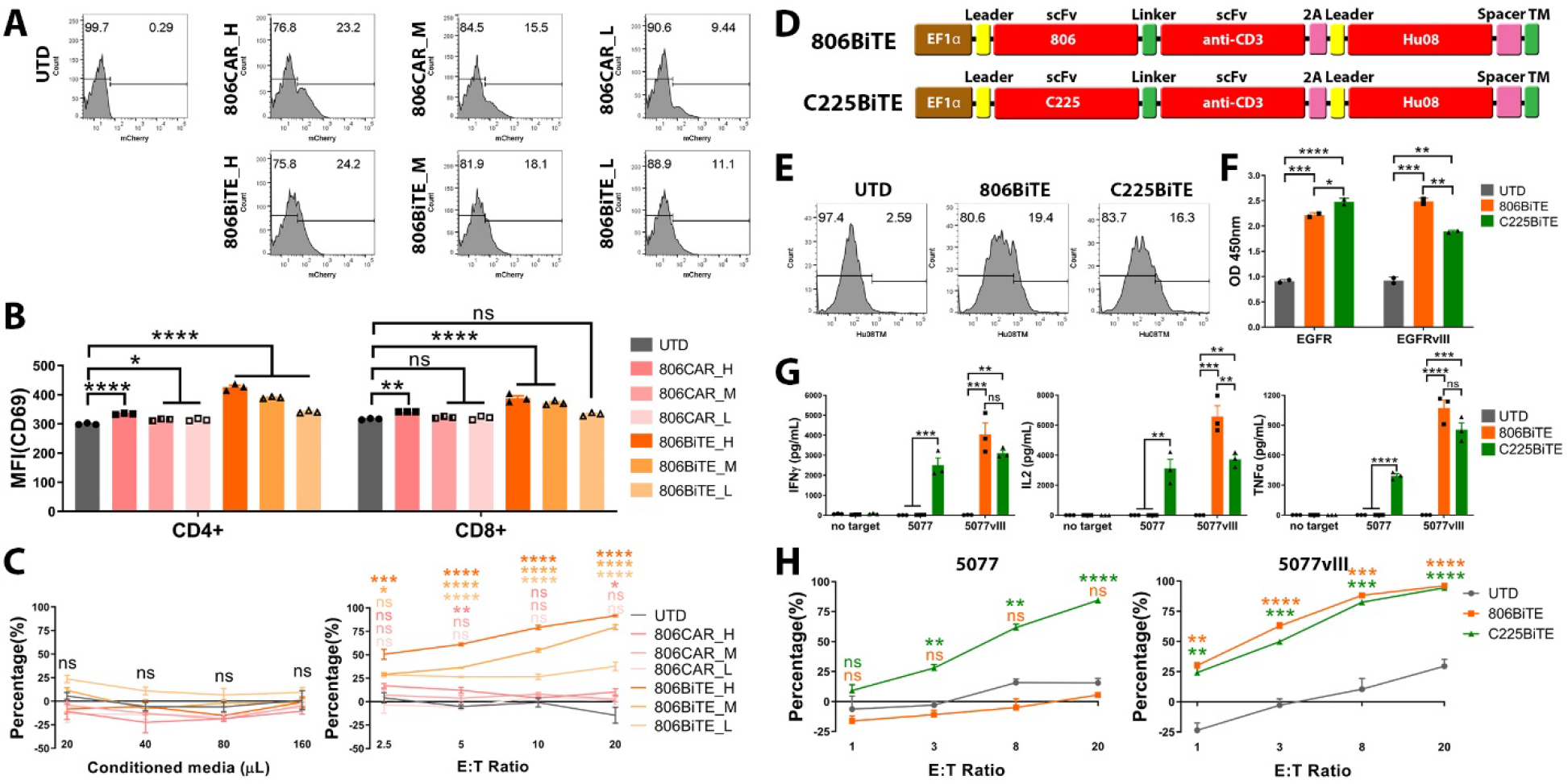
BiTE T cells respond to low antigen expression on glioma cancer stem cells. (A) Flow cytometry panels showing the high (H), medium (M), and low (L) levels of transduction of 806CARs (top row) and 806BiTEs (bottom row) by mCherry expression. (B) T cell activation, as measured by CD69 expression, in CD4+ and CD8+ T cells after 16hrs co-culture of 806CAR/BiTE T cells with 5077. (C) Bioluminescence cytotoxicity assays of UTD T cells co-cultured with 5077 target cells in the presence of conditioned media of 806CAR/BiTE T cells. The volume of conditioned media used in each co-culture was indicated along the x axis (left panel). Bioluminescence cytotoxicity assay of 806CAR/BiTE T cells co-cultured with 5077 at different effector/target (E:T) ratios (2.5:1, 5:1, 10:1, and 20:1) and compared with the UTD T cell group (right panel), showing an expression-dependent level of target killing by the 806BiTE T cells. (D) Vector maps of the 806 (top) and C225 (bottom) BiTE designs. (E) Flow panels showing the transduction of the BiTE constructs, as determined by Hu08TM staining. (F) Conditioned media of T cells in each group tested with a standard direct ELISA to evaluate the binding ability of secreted BiTEs to recombinant EGFR and EGFRvIII protein. (G) 806/C225BiTE T cells and UTD T cells were co-cultured with 5077 or 5077^EGFRvIII+^ cells. Cytokine secretion (IFNγ, IL2 and TNFα) was detected with ELISA. (H) Bioluminescence cytotoxicity assays of 806/C225BiTE T cells co-cultured with 5077 cells (left panel) and 5077^EGFRvIII+^ cells (right panel), analyzed at different effector/target (E:T) ratios (1:1, 3:1, 8:1, and 20:1) and compared with the UTD T cell group. Statistically significant differences were calculated by one-way ANOVA with post hoc Tukey test. ns, not significant; *p<0.05, **p<0.01, ***p<0.001, ****p<0.0001. Data are presented as means ± SEM.

To further demonstrate that the low level of endogenous EGFR was the cause for 806BiTE activity against 5077 cells, another EGFR targeting BiTE (C225BiTE) was generated (Figure 4D). C225’s epitope is found on the L2 domain of EGFR, which is exposed in both active and inactive states (*39*). Comparable transduction efficacy of 806BiTE T cells and C225BiTE T cells was achieved, as determined by staining of Hu08CAR lacking signaling motifs (Figure 4E). Conditioned media from 806BiTE and C225BiTE secreting T cells was collected after an overnight culture and used in a standard direct ELISA to confirm the secretion and binding of 806BiTE and C225BiTE to recombinant EGFR and EGFRvIII (Figure 4F). Secreted 806BiTEs and C225BiTEs significantly bound to both EGFR and EGFRvIII. EGFR binding was more prominent for C225BiTE-secreting T cells than for 806BiTE-secreting T cells (p=0.0273). Additionally, binding to EGFRvIII was more prominent for 806BiTE-secreting T cells than the C225BiTE-secreting T cells (p=0.0039). Cytokine secretion was detected in 806BiTE and C225BiTE T cells when co-cultured with either 5077 or 5077^EGFRvIII+^ cells. Significant cytokine secretion was detected in C225BiTE T cells co-cultured with 5077 cells when compared with UTD or 806BiTE T cell co-culture groups, indicating that C225BiTE-secreting T cells could be stimulated by the low levels of EGFR expressed in 5077. In co-cultures with 5077^EGFRvIII+^ cells, significant cytokine secretion was detected in both 806BiTE and C225BiTE T cell groups when compared with UTD T cells. IL-2 secretion was higher in the 806BiTE T cell group than in the C225BiTE T cell group, which is also indicated by the ELISA result of EGFRvIII binding by 806BiTE. Cytotoxicity was observed in C225BiTE T cells co-cultured with 5077 cells in a bioluminescence cytotoxicity assay, but not in UTD or 806BiTE T cell co-culture groups. In co-culture with 5077^EGFRvIII+^ cells, both 806BiTE and C225BiTE T cells showed potent cytotoxicity. Taken together, the low levels of EGFR found on 5077 glioma cells can be targeted by both 806BiTE-secreting T cells and C225BiTE-secreting T cells. However, this was not observed in 806CAR T cells or 806BiTE-recruited UTD T cells.

### 806BiTE consistent with the binding property of 806 antibody when responding to target cells

After determining the ability of 806BiTE T cells to respond to low levels of EGFR in GBM cells, it was necessary to examine the safety of 806BiTE T cells in responding to physiologic EGFR expressed on normal astrocytes (Figure 5A). To determine the response of 806BiTE T cells to astrocytes, we generated 806 BiTE T cells at high (H), medium (M), and low (L) levels of transduction efficacy and compared them with 806CAR T cells at comparable transduction levels as a control. We found T cell activation to be positively correlated with transduction levels after 806BiTE/CAR T cells were co-cultured with astrocytes overnight (Figure 5B). However, in contrast to the response to 5077 or 5077^EGFRvIII+^ glioma cells, the activation levels in 806BiTE T cell groups were significantly lower than that of 806CAR T cell groups at the same corresponding transduction levels, in both CD4+ and CD8+ T cells. Cytokine production was also detected in 806BiTE and 806CAR T cells after overnight co-culturing with astrocytes (Figure 5C). In 806CAR T cells, cytokine production was positively correlated with the transduction efficacy in both CD4+ and CD8+ T cells. Consistent with the results of T cell activation, cytokine production in 806BiTE T cell groups was significantly lower than 806CAR T cell groups in corresponding transduction efficacy in both CD4+ and CD8+ T cell subgroups. More strikingly, with the exception of TNFα production in CD4+ 806BiTE T cells in the high transduction efficacy subgroup, no statistical differences of cytokine production in any other 806BiTE T cell co-culture groups were observed when compared to the UTD T cell co-culture group.

**Figure 5.**
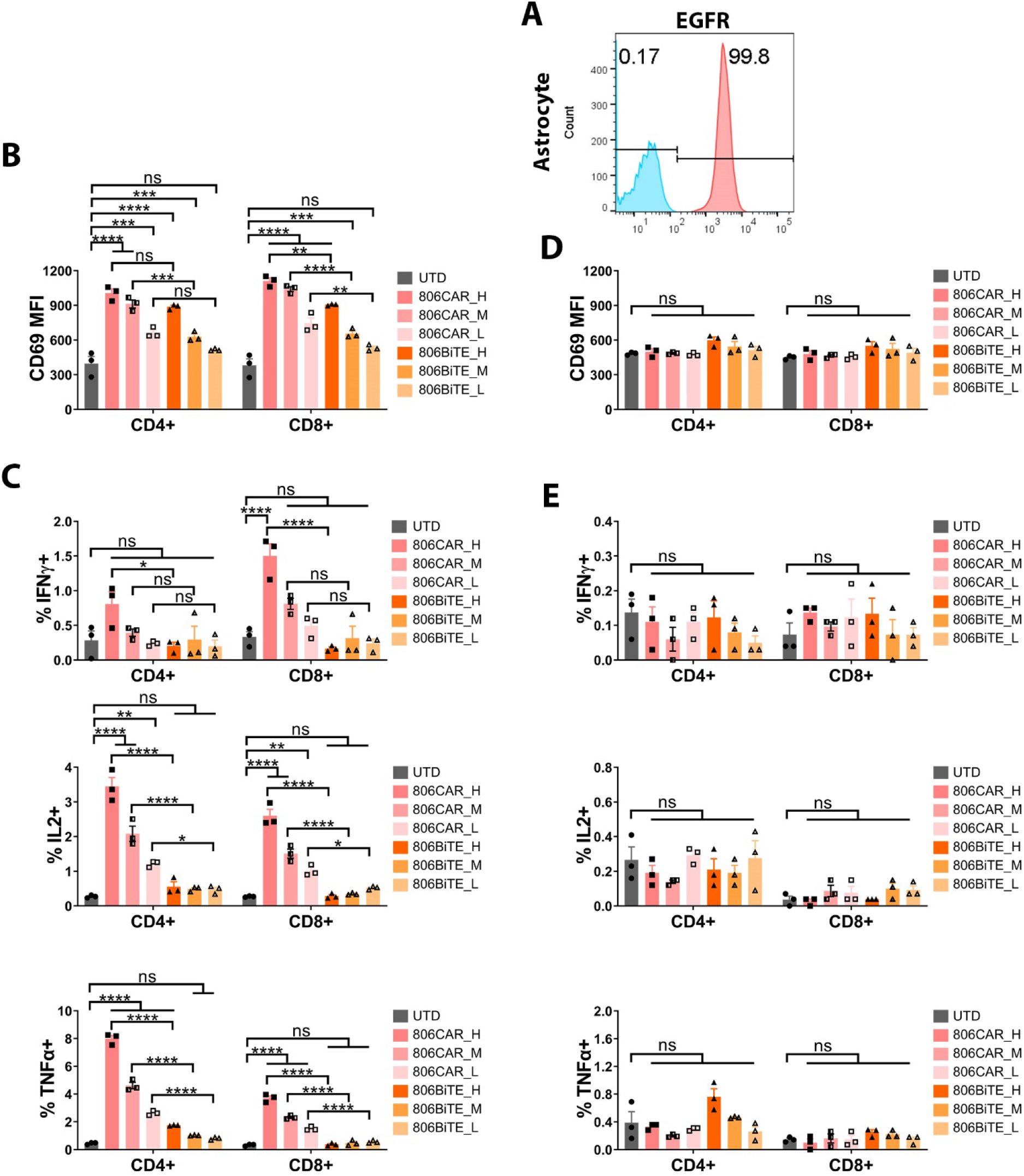
806BiTE shows comparable activity to 806CAR in co-culture with human astrocytes. A) Flow profile showing the expression of EGFR in astrocytes. (B) 806CAR/BiTE T cells with high (H), medium (M), and low (L) levels of transduction were co-cultured with astrocytes. T cell activation, as measured by CD69 expression, in CD4+ and CD8+ T cells after 16hrs co-culture. (C) Flow based intracellular cytokine (IFNγ, IL2 and TNFα) staining of 806CAR/BiTE T cells co-cultured with astrocytes. CD4 and CD8-positive subgroups of T cells were distinguished by human CD8 staining. (D) T cell activation, as demonstrated by CD69 expression, in CD4+ (top) and CD8+ (bottom) T cells after 16hrs co-culture with target cells and conditioned media of CAR/BiTE T cells. (E) Flow based intracellular cytokine (IFNγ, IL2 and TNFα) staining of UTD T cells co-cultured with astrocytes in conditioned media of CAR/BiTE T cells. Statistically significant differences were determined using log-rank test. ns, not significant; *p<0.05, **p<0.01, ***p<0.001, ****p<0.0001. Data are presented as means ± SEM.

To determine the ability of 806BiTE to engage UTD T cells and astrocytes, conditioned media of 806BiTE and 806CAR T cells at different levels of transduction efficacy was collected. CD69 expression and cytokine production was detected in UTD T cells co-cultured with astrocytes in the conditioned media from 806BiTE/806CAR T cells (Figure 5D and 5E). No significant upregulation of CD69 expression or cytokine production was detected in conditioned media from either 806BiTE or 806CAR T cell groups, when compared with conditioned media of UTD T cells. These results demonstrated that soluble 806BiTE and 806BiTEs secreted by T cells lead to weak activation of effector T cells in response to the EGFR expressed on astrocytes.

### BiTEs responded to target positive cells in bivalent targeting constructs

To deal with the heterogeneous antigen expression found on tumors, we generated gene-modified bivalent-targeting T cells simultaneously targeting tumor-expressed EGFR and IL13Rα2. To further assess the ability of BiTEs to be used in bivalent targeting, we generated 806BiTE-Hu08CAR, 806BiTE-Hu08BiTE T cells, and 806CAR-Hu08CAR T cells (Figure 6A). To mimic the ratio of transduced T cells with UTD T cells commonly seen in the infusion T cell products used in clinical trials (*18*), bivalent T cells with low transduction efficacy were adopted (Figure 6B). The human glioma line, D270, endogenously expresses both EGFR and IL13Rα2 and was used as target cells, in addition to 5077 cells (Figure 6C). All bivalent targeting T cells (806CAR-Hu08CAR, 806BiTE-Hu08CAR and 806BiTE-Hu08BiTE) demonstrated activation when co-cultured with target overexpressing 5077 or D270 cells (Figure 6D). Strikingly, there were differences in the activation of BiTE-producing T cells and CAR T cells in response to antigen-positive target cells. Both 806BiTE-Hu08CAR and 806BiTE-Hu08BiTE T cells showed superior response to 5077^EGFRvIII+^ cells and D270 cells when compared with 806CAR-Hu08CAR T cells. 806BiTE-Hu08BiTE T cells had a higher response to 5077^IL13Rα2+^ cells compared with 806CAR-Hu08CAR and 806BiTE-Hu08CAR T cells. Consistent results were also observed in cytokine production in both CD4+ and CD8+ subgroups (Figure 6E). We did not observe enhanced activation in the 806BiTE-Hu08BiTE population responding to D270 cells when compared to 806BiTE-Hu08CAR T cells. In cytotoxicity assays, all bivalent targeting T cells (806CAR-Hu08CAR, 806BiTE-Hu08CAR and 806BiTE-Hu08BiTE) showed dose-dependent cytotoxicity in target overexpressing 5077 and D270 cells (Figure 6F). Concordant with the above results, cytotoxicity of BiTE+ T cells responding to target antigen-positive cells was significantly higher than cytotoxicity observed in T cells expressing the equivalent CAR. No enhanced cytotoxicity was observed in 806BiTE-Hu08BiTE T cells responding to the double-positive D270 cells compared with 806BiTE-Hu08CAR T cells.

**Figure 6.**
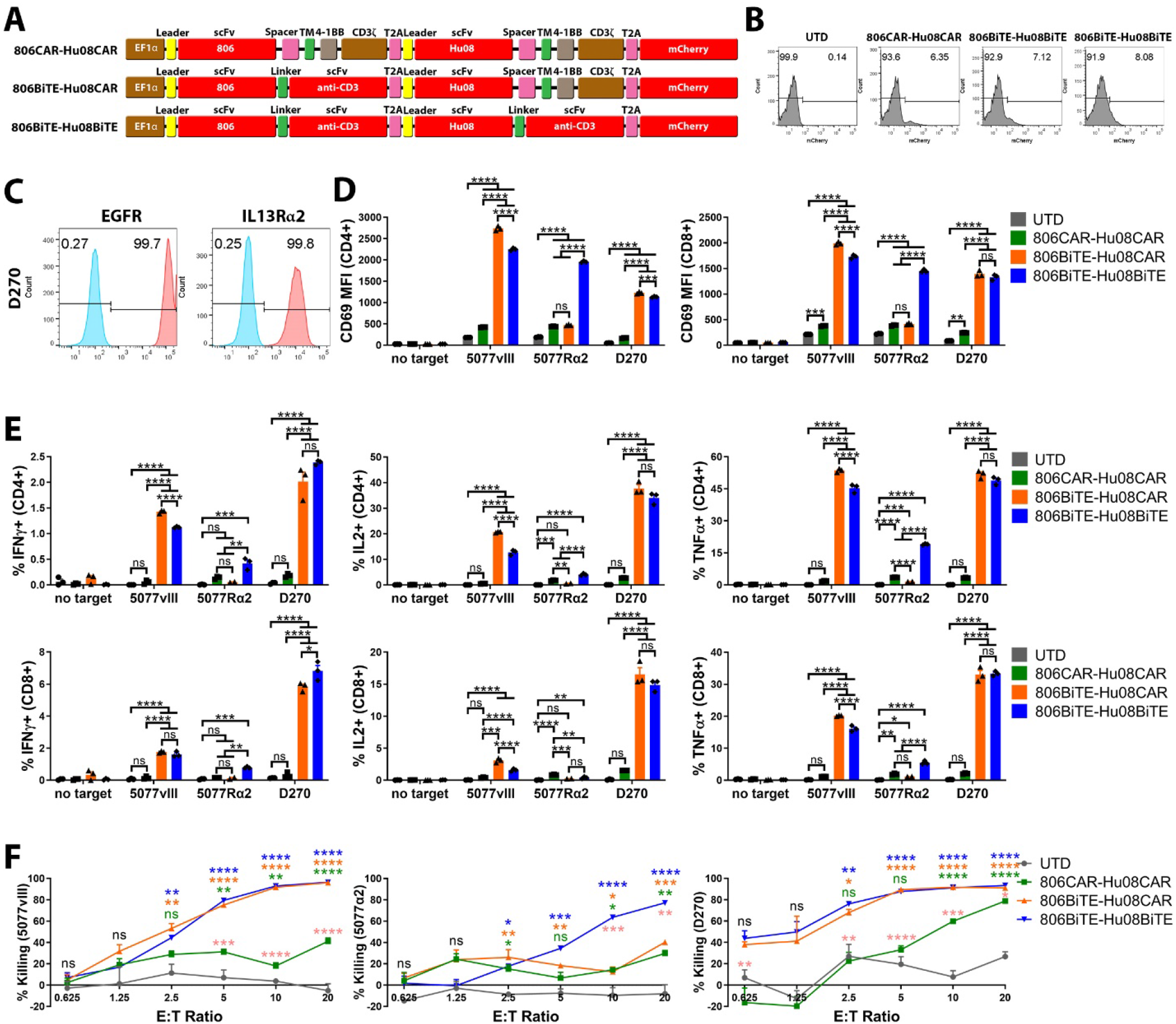
BiTE mediates anti-tumor activity in antigen positive cells in bivalent targeting constructs. (A) Vector maps of 806CAR-Hu08CAR, 806BiTE-Hu08CAR, and 806BiTE-Hu08BiTE constructs. (B) Flow cytometric detection of T cell transduction by mCherry expression. (C) Flow panel of EGFR (left) and IL13Rα2 (right) expression on glioma line D270. (D) Bivalent T cell activation, as measured by CD69 expression, in CD4+ (left) and CD8+ (right) T cells after 16hrs co-culture with D270. (E) Flow based intracellular cytokine (IFNγ, IL2 and TNFα) staining of bivalent targeting T cells co-cultured with target cells. CD4+ (top) and CD8+ (bottom) subgroups of T cells were distinguished by human CD8 staining. (F) Bioluminescence cytotoxicity assays of bivalent targeting T cells co-cultured with 5077^EGFRvIII+^ cells (left panel), 5077^IL13Rα2+^ cells (middle panel) and D270 cells (right panel), analyzed at different effector/target (E:T) ratios (0.625:1, 1.25:1, 2.5:1, 5:1, 10:1, and 20:1) and compared with the UTD T cell group. Statistically significant differences were calculated by one-way ANOVA with post hoc Tukey test. In cytotoxicity assays, pink asterisks indicate statistically differences between bivalent targeting T cell groups. ns, not significant; *p<0.05, **p<0.01, ***p<0.001, ****p<0.0001. Data are presented as means ± SEM.

To demonstrate the anti-tumor effects of bivalent BiTE targeting T cells *in vivo*, we first treated mice with 806BiTE-Hu08CAR T cells 8 days after D270 subcutaneous tumor cell implantation in NSG mice, using Hu08CAR T cells as a positive control (Figure 7A). Both 806BiTE-Hu08CAR and Hu08CAR T cells treated groups significantly controlled tumor growth and prolonged survival compared to mice treated with UTD T cells (p<0.0001). Notably, we observed earlier control of tumor growth in the 806BiTE-Hu08CAR cohort when compared to the Hu08CAR cohort. Tumor sizes were also significantly different in the 806BiTE-Hu08CAR T cell treated group versus the Hu08CAR T cell treated group on post-treatment day 6 (31.29mm^2^ versus 74.52mm^2^, p=0.0076) and day 9 (27.92mm^2^ versus 78.10mm^2^, p=0.0152). Next, we infused 806BiTE-Hu08CAR, 806BiTE-Hu08BiTE, and 806CAR-Hu08CAR T cells in D270 tumor bearing mice 8 days after subcutaneous tumor implantation (Figure 7B). All bivalent targeting T cells significantly controlled tumor growth (p<0.0001) and prolonged survival (p<0.001) as compared with the UTD T cell cohort. Tumor sizes in 806BiTE-Hu08CAR and 806BiTE-Hu08BiTE T cell groups significantly decreased on day 17 after tumor implantation (day 9 after T cells infusion), which was earlier than the 806CAR-Hu08CAR T cell infused group. Average tumor sizes in 806BiTE-Hu08CAR and 806BiTE-Hu08BiTE T cell groups were roughly 50% that of the 806CAR-Hu08CAR T cell group on day 14 and 17 after tumor implantation (day 6 and day 9 after T cells infusion), however no statistical difference was observed (41.84mm^2^ and 41.88mm^2^ versus 82.67mm^2^, p=0.0858 and 0.0863; 46.35mm^2^ and 43.83mm^2^ versus 86.50mm^2^, p=0.0882 and 0.0637). There was no difference in tumor size and survival between 806BiTE-Hu08CAR and 806BiTE-Hu08BiTE T cell groups. 806CAR-Hu08CAR T cells showed better control of tumor growth and mouse survival than 806BiTE-Hu08CAR and 806BiTE-Hu08BiTE T cell groups in the long run. However, the most potent anti-tumor control was found in bivalent BiTE secreting T cells (day 6 or day 9 after T cells treatment). Taken together, BiTE-secreting bivalent T cells demonstrated significant anti-tumor activity both *in vitro* and *in vivo*.

**Figure 7.**
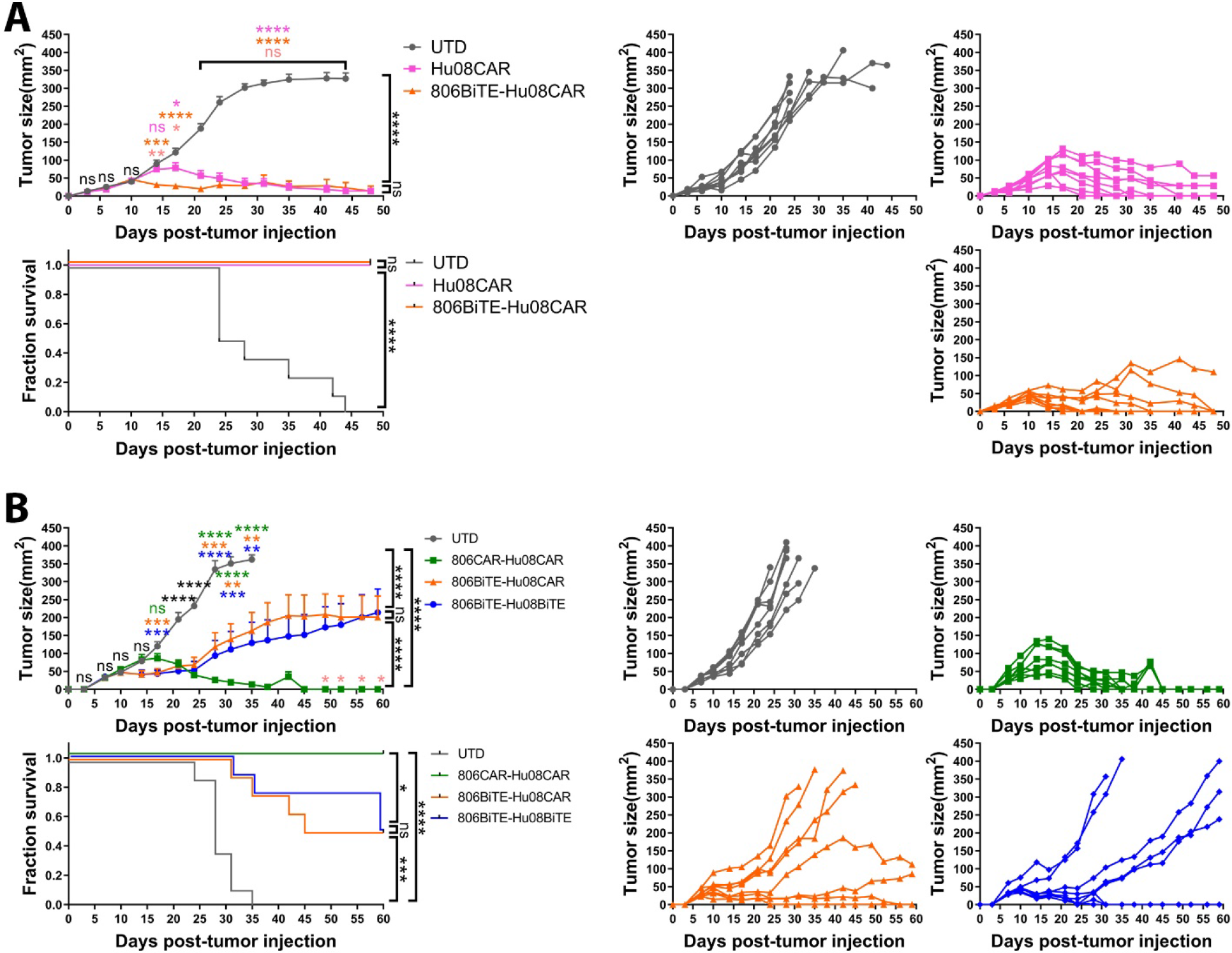
BiTE transduced T cells significantly delay early tumor growth in a GBM implanted mouse model. (A) 800,000 Hu08CAR and 806BiTE-Hu08CAR positive T cells or the same number of UTD T cells were injected intravenously eight days after D270 subcutaneous implantation. (B) 1,200,000 bivalent targeting construct-transduced T cells (806CAR-Hu08CAR, 806BiTE-Hu08CAR, and 806BiTE-Hu08BiTE) or the same number of UTD T cells were injected intravenously eight days after D270 subcutaneous implantation. Tumor size was compared between each group. Statistically significant differences of tumor size at each time point were calculated by one-way ANOVA with post hoc Tukey test. Pink asterisks indicate statistically significant differences between transduced T cell groups. Linear regression was used to test for significant differences between the experimental groups. Survival based on time to endpoint was plotted using a Kaplan-Meier curve (Prism software). Statistically significant differences were determined using log-rank test. ns, not significant; *p<0.05, **p<0.01, ***p<0.001, ****p<0.0001. Data are presented as means ± SEM.

## DISCUSSION

In this study, we generated T cell-secreting BiTEs directed to tumor-associated EGFR and IL13Rα2, both targets having previously been demonstrated to be expressed on GBM cells (*32, 40*). BiTEs secreted by T cells demonstrated superior anti-tumor activity when compared to their corresponding CAR version. 806BiTE-secreting T cells also demonstrated enhanced sensitivity in responding to low target antigen-expressing tumor cells. In bivalent targeting constructs, BiTEs also outperform CARs in killing target-positive tumor cells *in vitro* and *in vivo*.

BiTE therapy and CAR T cell therapy are two main approaches for redirecting T cells against tumors, both of which demonstrated promising effects in pre-clinical and clinical studies (*4, 7, 9, 19, 20*). To integrate the advantages of both strategies, T cells were engineered to constitutively secrete BiTEs to recruit bystander T cells to engage tumor cells (*12, 41–43*). We deemed it important to compare the anti-tumor activity of these two molecules in a head-to-head comparison. Consistent with reported results of mRNA electroporated BiTE T cells (*13*), lentivirally-transduced 806BiTE and Hu08BiTE T cells also demonstrated superior anti-tumor responses when compared to their corresponding 806CAR and Hu08CAR T cells. In our *in vivo* tumor models, mice treated with BiTE secreting T cells established tumor control earlier than groups not treated with BiTE secreting T cells. Remarkably, when co-cultured with EGFR-low 5077, only 806BiTE secreting T cells significantly attacked target cells, but not 806BiTE-recruited T cells or 806CAR T cells. Combined with a lack of cytotoxic activity against EGFR-expressing astrocytes (*32*), this sensitivity and selectivity highlights the safety features of 806BiTEs and 806BiTE secreting T cells. In a phase II clinical trial of blinatumomab treatment for refractory or relapsed leukemia, 43% patients achieved a complete response. This rate is raised to 82% in the treatment of patients with minimal residual disease (*8*), which could be attributed to the remarkable sensitivity of BiTEs in responding to tumor cells (*44*). In our study, the sensitivity of the T cell secreted 806BiTE was consistently higher than that of 806CAR T cells, again suggesting that BiTE secreting cells is a viable strategy in the treatment of antigen-low tumors or with heterogeneous antigen expression in tumors.

Compared with monovalent CAR T cells or bivalent CAR T cells, BiTE-CAR T cells and BiTE-BiTE T cells showed superior control of tumor growth in a glioma mouse model 9 days after T cell treatment. 20 days after T cell administration, BiTE-CAR T cells and BiTE-BiTE T cells demonstrated loss of tumor control, but this phenomenon was not found in mice treated with bivalent CAR T cells. One potential explanation for the lack of long-term control by BiTEs is the limited pool of unmodified human bystander T cells available for a BiTE to recruit in the immunodeficient NSG mouse system. Unmodified T cells may not persist in an NSG mouse 20 days after T cell administration, given the lack of stimulation and engraftment (*45*). Additionally, BiTE-mediated T cell activation does not provide a co-stimulatory signal, presumably resulting in T-cell exhaustion and an inability to proliferate (*24*). Co-stimulatory signals such as 4-1BB and CD28 have significantly enhanced BiTE mediated T cell activation (*46, 47*). These reports suggest that providing an additional co-stimulatory signal in the BiTEs would be a promising strategy in maintaining the function of BiTE-secreting T cells and integrating the advantages of BiTE and CAR T cell therapy. Based on our results, repeated infusions of BiTE-secreting T cells could be a potent way to control tumor growth and provide long-term anti-tumor activity.

Antigen loss is a primary mechanism of resistance to redirected T cell therapies in cancer (*14*). CD19 is homogeneously expressed in mature B cells. Even so, approximately 7-25% of patients relapsed with CD19-negative disease after treatment with CD19-specific CAR T cells (*1*). In solid tumors, heterogeneous antigen expression is much more common. In a single-cell RNA-seq study, 430 cells from five GBMs resulted in distinct expression patterns among individual tumor cells derived from the same tumor (*28*). A transient complete response was achieved in a patient treated with IL13Rα2-specific CAR T cells despite the GBM having heterogeneous IL13Rα2 expression. However, the patient’s tumor eventually recurred with target antigen loss (*6*). In our trial of CAR T cells redirected to EGFRvIII, loss of the target antigen was observed in the majority of patients with surgical resection post-CAR T infusion. However, loss of EGFRvIII did not have a significant influence on wild-type amplified EGFR expression in these tumors (*18*). A multivalent targeting strategy was demonstrated as a valid way in mitigating antigen loss to treat hematological and solid tumors (*15, 17, 21–23*). In this study, we also detected heterogeneously expressed EGFRvIII and EGFR in freshly resected glioma samples. EGFR was found to be more prominent than EGFRvIII expression in individual tumor cells, indicating that perhaps a meaningful clinical outcome can be achieved by targeting amplified, oncogenic wild-type EGFR instead of EGFRvIII.

A previous study confirmed that EGFRvIII CAR T cells can traffic to the site of EGFRvIII positive tumors and secrete EGFR BiTEs into the microenvironment (*24*). Extending this observation, we adopted the EGFR conformation-specific 806 scFv to generate an 806BiTE. This construct is another strategy to target both EGFR and EGFRvIII with a single scFv. We also found IL13Rα2 to often be co-expressed in the same tumor with EGFR and EGFRvIII. Based on the TCGA data, both IL13Rα2 and EGFR expression positively correlated with WHO glioma grade. EGFRvIII was also reported to promote IL13Rα2 mediated tumor proliferation by directly binding its intracellular signaling motif (*29*). Simultaneous targeting of IL13Rα2, EGFR, and EGFRvIII by T cells could result in meaningful clinical outcomes in the treatment of gliomas. In bivalent targeting constructs, BiTEs also mediated superior target specific responses compared to CAR T cells and established earlier control of tumor growth. Besides superior sensitivity to low target antigen expression, bivalent targeting using BiTEs demonstrated potency in mitigating the problem of tumor antigen heterogeneity.

806 binds to the CR1 domain of EGFR, which is masked in the inactive monomer state. Exposure of this epitope preferentially occurs under tumor-specific conditions, such as constitutive EGFR activation via EGFR amplification and EGFRvIII mutations (*34, 48*). This conformational specificity makes 806 bind to tumor associated oncogenic EGFR and EGFRvIII. Our group has previously confirmed 806CAR T cells successfully target other EGFR oncogenic mutations, including extracellular missense mutants EGFR^A289V^ and EGFR^R108K^, while demonstrating little cytotoxicity against human astrocytes (*32*). Similarly, there was no overt response to astrocytes with 806BiTE T cells or 806BiTE-recruited UTD T cells. Given these results, BiTE-redirected T cell therapies could be a tumor-specific approach when utilizing a confirmation-specific scFv. 806BiTE T cells also demonstrated a more potent response to EGFRvIII than C225BiTE T cells. C225BiTE T cells did respond to the EGFR present on 5077 cells, while 806BiTE T cells did not show a response. This result may possibly be explained by the different binding affinity of 806 and C225 scFvs with EGFRvIII and EGFR (*39, 49*), which is also concordant with our results of ELISA-based BiTE binding assays. In our study, BiTE binding on T cells was only detected in the presence of target antigen. That may be attributed to the low affinity of the anti-CD3 scFv (*50*). Consistent with a previous study (*51*), we did not detect BiTE mediated T cell activation in the absence of target antigen.

In summary, BiTE T cells demonstrated superior activation, sensitivity, and specificity in responding to their cognate antigen than the corresponding CAR construct. BiTE T cells and BiTE-CAR T cells are promising strategies in addressing the clinical problem of targeting solid tumor antigen heterogeneity and may result in paving the way for targeted T cell therapies.

## MATERIALS AND METHODS

### Cell Lines and Culture

Human GSC lines (5041, 5077, 5391, 4701, 4860, 5377, 5560, 4806 4957 and 4892) were isolated from patient excised tumor tissue (Department of Neurosurgery, Perelman School of Medicine, Philadelphia, PA) and maintained in Dulbecco’s Modified Eagle’s Medium/Nutrient Mixture F12 Ham with penicillin/streptomycin, GlutaMAX-1, B27 minus A, epidermal growth factor and basic fibroblast growth factor (Corning, Corning, NY). EGFR copy number amplifications and EGFRvIII detection was carried out as previously described, using the Center for Personalized Diagnostics at the University of Pennsylvania (*36*). GSC line 5077 was lentivirally transduced to express or co-express EGFRvIII, IL13Rα2, click beetle green luciferase and green fluorescent protein (GFP) under control of the EF-1α promoter. D270 glioma cells were grown and passaged in the right flanks of NSG mice to keep their glioma characteristics *in vivo*. Human astrocytes were purchased from ScienCell Research Laboratories (Carlsbad, CA) and maintained in culture for 3 to 7 passages in medium, as directed by the vendor.

### Vector constructs

The nucleic acid sequences of EGFR targeting scFv (806) or IL13Rα2 targeting scFv (Hu08) in a second-generation CAR construct or a BiTE construct were synthesized and ligated into pTRPE lentiviral vector with a T2A ribosomal skipping sequence and an mCherry gene (Twist Bioscience, San Francisco, CA). 806 CAR sequences and 806 BiTE sequences with P2A ribosomal skipping sequences were synthesized, digested with NheI and XbaI and ligated into pTRPE vector of an Hu08BBz T2A mCherry construct to generate bivalent targeting constructs. Truncated Hu08 CAR sequences (Hu08TM; Hu08 scFv with leader, hinge, and transmembrane sequences of human CD8α) were digested with XbaI and SalI and ligated into the 806BiTE and Hu08BBz bivalent targeting structure in the same enzyme sites to replace Hu08BBz and mCherry genes. Hu08TM was used as a cell surface tag. C225 BiTE structure was digested with NheI and HpaI to replace the 806 BiTE in the Hu08TM tag construct.

### Human T Cell Transduction and Culture *In Vitro*

Human T cells transduction and culture were performed as previously described (*52*). Briefly, isolated T cells were derived from leukapheresis products obtained from the Human Immunology Core at the University of Pennsylvania, using de-identified healthy donors under an institutional review board-approved protocol. T cells were stimulated with Dynabeads Human T-Activator CD3/CD28 (Life Technologies, Carlsbad, CA) at a bead-to-cell ratio of 3:1. After 24-hr stimulation, lentivirus was added into the culture media and thoroughly mixed to produce stably transduced CAR T cells. The concentration of the expanding human T cells was calculated on a Coulter Multisizer (Beckman Coulter, Brea, CA) and maintained at 1.0-2.0 × 10^6^ cells per mL in R10 media (RPMI-1640 plus GlutaMAX-1, HEPES, pyruvate and penicillin/streptomycin (Thermo Fisher Scientific), supplemented with 10% fetal bovine serum (FBS)) and 30IU/mL recombinant human IL-2 (rhIL-2; Thermo Fisher Scientific, Waltham, MA).

### Flow Cytometry

Fresh human GBM samples were minced, and single cell suspensions were washed through a cell strainer (40μm). Red blood cells were lysed with Ammonium-Chloride-Potassium (ACK) Lysing Buffer (Lonza, Basel, Switzerland). The size and concentration of cells was measured on a Coulter Multisizer after washing with PBS. GBM cells were distinguished with live/dead viability stain (Thermo Fisher Scientific), followed by human CD45 (BioLegend, San Diego, CA) stain. Cell lines were digested from culture *in vitro* and stained with live/dead viability stain, BV711/PE conjugated anti-IL13Rα2 (BioLegend), PE conjugated anti-EGFR (BioLegend) and non-conjugated anti-EGFRvIII antibody (Novartis, Basil, Switzerland). PE conjugated anti-Rabbit IgG (BioLegend) secondary stains were used for detecting these targets. mCherry and Hu08TM were used as tags of T cell transduction. Cells expressing Hu08TM were stained with biotinylated protein L (GenScript, Piscataway, NJ) and secondary detection was achieved by the addition of streptavidin-coupled PE (BD Biosciences, Franklin Lakes, NJ).

In co-culture experiments, transduced or UTD T cells (2×10^5^ cells per well in 100μL R10 media) were co-cultured with different target cells (2×10^5^ cells per well in 100μL R10 media) in 96-well round bottom tissue culture plates, at 37°C with 5% CO_2_ for 16hrs. The supernatant of the T cells was used to stimulate UTD T cells co-cultured with target cells. Transduced or UTD T cells (4×10^5^ cells in 200μL R10 media) were cultured in 96-well round bottom tissue culture plates at 37°C in 5% CO_2_ for 16hrs. 160μL supernatant was taken from each well and added into UTD T cells (2×10^5^ cells) co-cultured with target cells (2×10^5^ cells) to reach 200μL per well. Human CD4+ and CD8+ T cells were distinguished with live/dead viability stain (Thermo Fisher Scientific), followed by human CD3 and CD8 (BioLegend) stain. Biotinylated protein L (GenScript) and the addition of streptavidin-coupled PE (BD Biosciences) was used to detect BiTEs’ binding to T cells. APC conjugated anti-human CD69 (BioLegend) was used to detect the T cell stimulation. Before and after each staining, cells were washed twice with PBS containing 2% fetal bovine serum (FACS buffer). Fluorescence was assessed using a BD LSRFortessa flow cytometer and data were analyzed with FlowJo software.

### Immunohistochemistry

Formalin-fixed paraffin-embedded tissue sections, 5μm thick, were stained using antibody against IL13Rα2 (clone E7U7B Cell Signaling 85677, dilution 1:100, Danvers, MA) and EGFRVIII (clone D6T2Q Cell Signaling 64952S, dilution 1:100). The double staining was done sequentially on a Leica Bond-IIITM instrument using the Bond Polymer Refine DAB Detection System (Leica Microsystems DS9800, Buffalo Grove, IL) and Refine Red Detection System (Leica Microsystems DS9390). Heat-induced epitope retrieval was done in ER2 solution (Leica Bio systems AR9640) for 20 minutes. The entire experiment was done at room temperature. Slides were washed three times between each step with bond wash buffer or water.

### Intracellular Cytokine Analysis

To stimulate UTD T cells, supernatants of T cells were collected in 96-well round bottom tissue culture plates as described above and added into UTD T cells (2×10^5^ cells) co-cultured with target cells (2×10^5^ cells) in 96-well round bottom tissue culture plates in R10 media with the presence of Golgi inhibitors monensin and brefeldin A (BD Bioscience) to reach 200μL per well. When transduced T cells were used to respond to target cells, transduced or UTD T cells (2×10^5^ cells in 160μL R10 media) were cultured in 96-well round bottom tissue culture plates at 37℃ in 5% CO_2_ for 16hrs. Target cells (2×10^5^ cells in 40μL) with Golgi inhibitors were added into each well. 16hrs later, cells were washed, stained with live/dead viability stain, followed by surface staining for human CD3 and CD8 (BioLegend), then fixed and permeabilized, and intracellularly stained for human IFNγ, IL2 and TNFα. Cells were analyzed by flow cytometry (BD LSRFortessa) and gated on live, single-cell lymphocytes and CD3-positive lymphocytes.

### Bioluminescence Cytotoxicity Assay

1×10^4^ target cells transduced with click beetle green luciferase were seeded in 96-well plates with opaque walls. In the assays of supernatants, 5×10^4^ UTD T cells were added in each well. Supernatants of specific volumes (160μL, 80μL, 40μL, 20μL, et al) plus R10 media were added into each well to reach 200μL in total. No supernatant was added in control wells. In the assays of transduced T cells, effector cells were added at different E:T ratios, as indicated in each figure, to reach 200μL per well. No T cells were added in control wells. Plates were incubated at 37°C in 5% CO_2_ in an incubator. At the end of the incubation period, 15μg D-Luciferin (Gold Biotechnology, St. Louis, MO) in 5μL PBS was added in each well and incubated at room temperature for 10mins. Luminescence from the assay plates was measured with BioTek Synergy H4 hybrid multi-model microplate reader. Percentage specific lysis was calculated as follows: (control counts – sample counts)/control counts × 100%.

### Enzyme-linked immunosorbent assay (ELISA)

In order to detect the binding of BiTEs to their targets, a standard direct ELISA was performed with DuoSet Ancillary Reagent Kit 2 (R&D systems, Minneapolis, MN). After coating wells with recombinant human EGFR (1μg/mL), EGFRvIII (1μg/mL) and IL13Rα2 (4μg/mL) protein (Sino Biological, Wayne, PA), a 96-well plate was loaded with supernatants which were collected as described above, followed by biotin conjugated goat anti-mouse IgG (Jackson ImmunoResearch, West Grove, PA) detection antibody and secondary detection by the addition of horseradish peroxidase conjugated streptavidin. For detecting IFNγ, IL2 and TNFα by ELISA, supernatant was collected from T cells and target cells after 16hrs co-culture at a 1:1 ratio. The detection was performed with DuoSet ELISA kits (R&D Systems) as the introduction indicated.

### Mouse Model

All mouse experiments were conducted according to Institutional Animal Care and Use Committee (IACUC)-approved protocols and as previously described (*52*). NSG mice were injected with 5×10^5^ D270 tumor cells subcutaneously in 100μL of PBS on day 0. Tumor size was measured by calipers in two dimensions, L×W, for the duration of the experiment. T cells were injected in 100μL of PBS intravenously via the tail vein 8 days after tumor implantation. Survival was followed over time until a predetermined IACUC-approved endpoint was reached.

### Statistical Analysis

Data are presented as means ± SEM. Target detection on fresh tissue was analyzed with two-tail paired student t test. RNA-seq data from TCGA was analyzed with Kruskal-Wallis test. Survival curves were analyzed with Kaplan-Meier (log-rank test). The remaining experiments were analyzed with one-way Analysis of Variance (ANOVA) with post hoc Tukey test to compare the differences in each group. For the *in vivo* tumor study, linear regression was used to test for significant differences between the experimental groups. Survival, based on time to experimental endpoint, was plotted using a Kaplan-Meier curve. All statistical analyses were performed with Prism software version 9.0 (GraphPad, La Jolla, CA).

## ACKNOWLEDGEMENTS

The authors thank the Human Immunology Core at the University of Pennsylvania for providing T cells for the described work, the Stem Cell and Xenograft Core at the University of Pennsylvania for assistance with the animal work, the Small Animal Imaging Facility at the University of Pennsylvania for the bioluminescence imaging, and the Pathology Clinical Service Center of Penn Medicine for immunohistochemistry staining of IL13Rα2 and EGFRvIII, as well as Laura A. Johnson, Donald L. Siegel and Avery D. Posey, Jr. for the technical support on molecular biology.

## AUTHOR CONTRIBUTIONS

D.M.O., Z.A.B., Y.Y. designed the experiments. Y.Y., J.L.R., N.L., R.T., M.P.N, L.H., L.Z., J.V.Z, D.K., L.H., E.X.J., Y.L., H.L., J.Q., X.Z., L.Z., Z.A.B., D.M.O. performed the experiments. D.M.O., Z.A.B, Z.L., Y.Y. provided funding. All the authors contributed to the writing and editing of the manuscript.

## REFERENCES

1. S. Rafiq, C. S. Hackett, R. J. Brentjens, Engineering strategies to overcome the current roadblocks in CAR T cell therapy. Nat Rev Clin Oncol 17, 147–167 (2020).

2. S. L. Maude et al., Chimeric antigen receptor T cells for sustained remissions in leukemia. N. Engl. J. Med. 371, 1507–1517 (2014).

3. S. J. Schuster et al., Chimeric Antigen Receptor T Cells in Refractory B-Cell Lymphomas. N Engl J Med 377, 2545–2554 (2017).

4. E. A. Chong, M. Ruella, S. J. Schuster, P. Lymphoma Program Investigators at the University of, Five-Year Outcomes for Refractory B-Cell Lymphomas with CAR T-Cell Therapy. N Engl J Med 384, 673–674 (2021).

5. N. Raje et al., Anti-BCMA CAR T-Cell Therapy bb2121 in Relapsed or Refractory Multiple Myeloma. N Engl J Med 380, 1726–1737 (2019).

6. C. E. Brown et al., Regression of Glioblastoma after Chimeric Antigen Receptor T-Cell Therapy. N. Engl. J. Med. 375, 2561–2569 (2016).

7. M. E. Goebeler, R. C. Bargou, T cell-engaging therapies – BiTEs and beyond. Nat Rev Clin Oncol 17, 418–434 (2020).

8. M. S. Topp et al., Safety and activity of blinatumomab for adult patients with relapsed or refractory B-precursor acute lymphoblastic leukaemia: a multicentre, single-arm, phase 2 study. Lancet Oncol 16, 57–66 (2015).

9. H. Kantarjian et al., Blinatumomab versus Chemotherapy for Advanced Acute Lymphoblastic Leukemia. N Engl J Med 376, 836–847 (2017).

10. M. E. Goebeler et al., Bispecific T-Cell Engager (BiTE) Antibody Construct Blinatumomab for the Treatment of Patients With Relapsed/Refractory Non-Hodgkin Lymphoma: Final Results From a Phase I Study. J Clin Oncol 34, 1104–1111 (2016).

11. V. Dufner et al., Long-term outcome of patients with relapsed/refractory B-cell non-Hodgkin lymphoma treated with blinatumomab. Blood Adv 3, 2491–2498 (2019).

12. K. Iwahori et al., Engager T cells: a new class of antigen-specific T cells that redirect bystander T cells. Mol Ther 23, 171–178 (2015).

13. X. Liu et al., Improved anti-leukemia activities of adoptively transferred T cells expressing bispecific T-cell engager in mice. Blood Cancer J 6, e430 (2016).

14. R. G. Majzner, C. L. Mackall, Tumor Antigen Escape from CAR T-cell Therapy. Cancer Discov 8, 1219–1226 (2018).

15. T. J. Fry et al., CD22-targeted CAR T cells induce remission in B-ALL that is naive or resistant to CD19-targeted CAR immunotherapy. Nat Med 24, 20–28 (2018).

16. M. Hamieh et al., CAR T cell trogocytosis and cooperative killing regulate tumour antigen escape. Nature 568, 112–116 (2019).

17. D. Schneider et al., Trispecific CD19-CD20-CD22-targeting duoCAR-T cells eliminate antigen-heterogeneous B cell tumors in preclinical models. Sci Transl Med 13, (2021).

18. D. M. O’Rourke et al., A single dose of peripherally infused EGFRvIII-directed CAR T cells mediates antigen loss and induces adaptive resistance in patients with recurrent glioblastoma. Sci Transl Med 9, (2017).

19. A. Sternjak et al., Preclinical Assessment of AMG 596, a Bispecific T-cell Engager (BiTE) Immunotherapy Targeting the Tumor-specific Antigen EGFRvIII. Mol. Cancer Ther. 20, 925–933 (2021).

20. K. C. Pituch et al., Neural stem cells secreting bispecific T cell engager to induce selective antiglioma activity. Proc. Natl. Acad. Sci. U.S.A. 118, (2021).

21. K. Bielamowicz et al., Trivalent CAR T cells overcome interpatient antigenic variability in glioblastoma. Neuro Oncol. 20, 506–518 (2018).

22. M. Hegde et al., Combinational targeting offsets antigen escape and enhances effector functions of adoptively transferred T cells in glioblastoma. Mol Ther 21, 2087–2101 (2013).

23. M. Hegde et al., Tandem CAR T cells targeting HER2 and IL13Ralpha2 mitigate tumor antigen escape. J Clin Invest 126, 3036–3052 (2016).

24. B. D. Choi et al., CAR-T cells secreting BiTEs circumvent antigen escape without detectable toxicity. Nat Biotechnol 37, 1049–1058 (2019).

25. A. Omuro, L. M. DeAngelis, Glioblastoma and other malignant gliomas: a clinical review. JAMA 310, 1842–1850 (2013).

26. P. Y. Wen, S. Kesari, Malignant gliomas in adults. N Engl J Med 359, 492–507 (2008).

27. R. Stupp et al., Maintenance Therapy With Tumor-Treating Fields Plus Temozolomide vs Temozolomide Alone for Glioblastoma: A Randomized Clinical Trial. JAMA 314, 2535–2543 (2015).

28. A. P. Patel et al., Single-cell RNA-seq highlights intratumoral heterogeneity in primary glioblastoma. Science 344, 1396–1401 (2014).

29. J. P. Newman et al., Interleukin-13 receptor alpha 2 cooperates with EGFRvIII signaling to promote glioblastoma multiforme. Nat. Commun. 8, 1913 (2017).

30. Y. Yin et al., Checkpoint Blockade Reverses Anergy in IL-13Ralpha2 Humanized scFv-Based CAR T Cells to Treat Murine and Canine Gliomas. Mol Ther Oncolytics 11, 20–38 (2018).

31. A. C. Ravanpay et al., EGFR806-CAR T cells selectively target a tumor-restricted EGFR epitope in glioblastoma. Oncotarget 10, 7080–7095 (2019).

32. R. Thokala et al., High Affinity Chimeric Antigen Receptor with Cross-Reactive scFv to Clinically Relevant EGFR Oncogenic Isoforms. bioRxiv, 2021.2002.2004.429797 (2021).

33. L. Orellana et al., Oncogenic mutations at the EGFR ectodomain structurally converge to remove a steric hindrance on a kinase-coupled cryptic epitope. Proc Natl Acad Sci U S A 116, 10009–10018 (2019).

34. T. P. Garrett et al., Antibodies specifically targeting a locally misfolded region of tumor associated EGFR. Proc Natl Acad Sci U S A 106, 5082–5087 (2009).

35. T. G. Johns et al., The antitumor monoclonal antibody 806 recognizes a high-mannose form of the EGF receptor that reaches the cell surface when cells over-express the receptor. FASEB J 19, 780–782 (2005).

36. M. P. Nasrallah et al., Molecular Neuropathology in Practice: Clinical Profiling and Integrative Analysis of Molecular Alterations in Glioblastoma. Provisional Acceptance at Academic Pathology, (2019).

37. R. G. Verhaak et al., Integrated genomic analysis identifies clinically relevant subtypes of glioblastoma characterized by abnormalities in PDGFRA, IDH1, EGFR, and NF1. Cancer Cell 17, 98–110 (2010).

38. L. Roccograndi et al., SHP2 regulates proliferation and tumorigenicity of glioma stem cells. J. Neurooncol., (2017).

39. S. Li et al., Structural basis for inhibition of the epidermal growth factor receptor by cetuximab. Cancer Cell 7, 301–311 (2005).

40. Y. Yin et al., Checkpoint blockade reverses anergy in IL13Rα2 humanized scFv based CAR T cells to treat murine and canine gliomas. Molecular Therapy – Oncolytics, (2018).

41. B. Blanco, M. Compte, S. Lykkemark, L. Sanz, L. Alvarez-Vallina, T Cell-Redirecting Strategies to ‘STAb’ Tumors: Beyond CARs and Bispecific Antibodies. Trends Immunol 40, 243–257 (2019).

42. C. L. Bonifant et al., CD123-Engager T Cells as a Novel Immunotherapeutic for Acute Myeloid Leukemia. Mol Ther 24, 1615–1626 (2016).

43. M. P. Velasquez et al., T cells expressing CD19-specific Engager Molecules for the Immunotherapy of CD19-positive Malignancies. Sci Rep 6, 27130 (2016).

44. S. Offner, R. Hofmeister, A. Romaniuk, P. Kufer, P. A. Baeuerle, Induction of regular cytolytic T cell synapses by bispecific single-chain antibody constructs on MHC class I-negative tumor cells. Mol Immunol 43, 763–771 (2006).

45. P. S. Hegde, D. S. Chen, Top 10 Challenges in Cancer Immunotherapy. Immunity 52, 17–35 (2020).

46. C. Claus et al., Tumor-targeted 4-1BB agonists for combination with T cell bispecific antibodies as off-the-shelf therapy. Sci Transl Med 11, (2019).

47. C. E. Correnti et al., Simultaneous multiple interaction T-cell engaging (SMITE) bispecific antibodies overcome bispecific T-cell engager (BiTE) resistance via CD28 co-stimulation. Leukemia 32, 1239–1243 (2018).

48. H. K. Gan, A. W. Burgess, A. H. Clayton, A. M. Scott, Targeting of a conformationally exposed, tumor-specific epitope of EGFR as a strategy for cancer therapy. Cancer Res 72, 2924–2930 (2012).

49. T. G. Johns et al., Identification of the epitope for the epidermal growth factor receptor-specific monoclonal antibody 806 reveals that it preferentially recognizes an untethered form of the receptor. J. Biol. Chem. 279, 30375–30384 (2004).

50. T. Dreier et al., Extremely potent, rapid and costimulation-independent cytotoxic T-cell response against lymphoma cells catalyzed by a single-chain bispecific antibody. Int J Cancer 100, 690–697 (2002).

51. K. Brischwein et al., Strictly target cell-dependent activation of T cells by bispecific single-chain antibody constructs of the BiTE class. J Immunother 30, 798–807 (2007).

52. L. A. Johnson et al., Rational development and characterization of humanized anti-EGFR variant III chimeric antigen receptor T cells for glioblastoma. Sci. Transl. Med. 7, 275ra222 (2015).

