## Supplementary Figure 1 for "Locally secreted BiTEs complement CAR T cells by enhancing solid tumor killing"

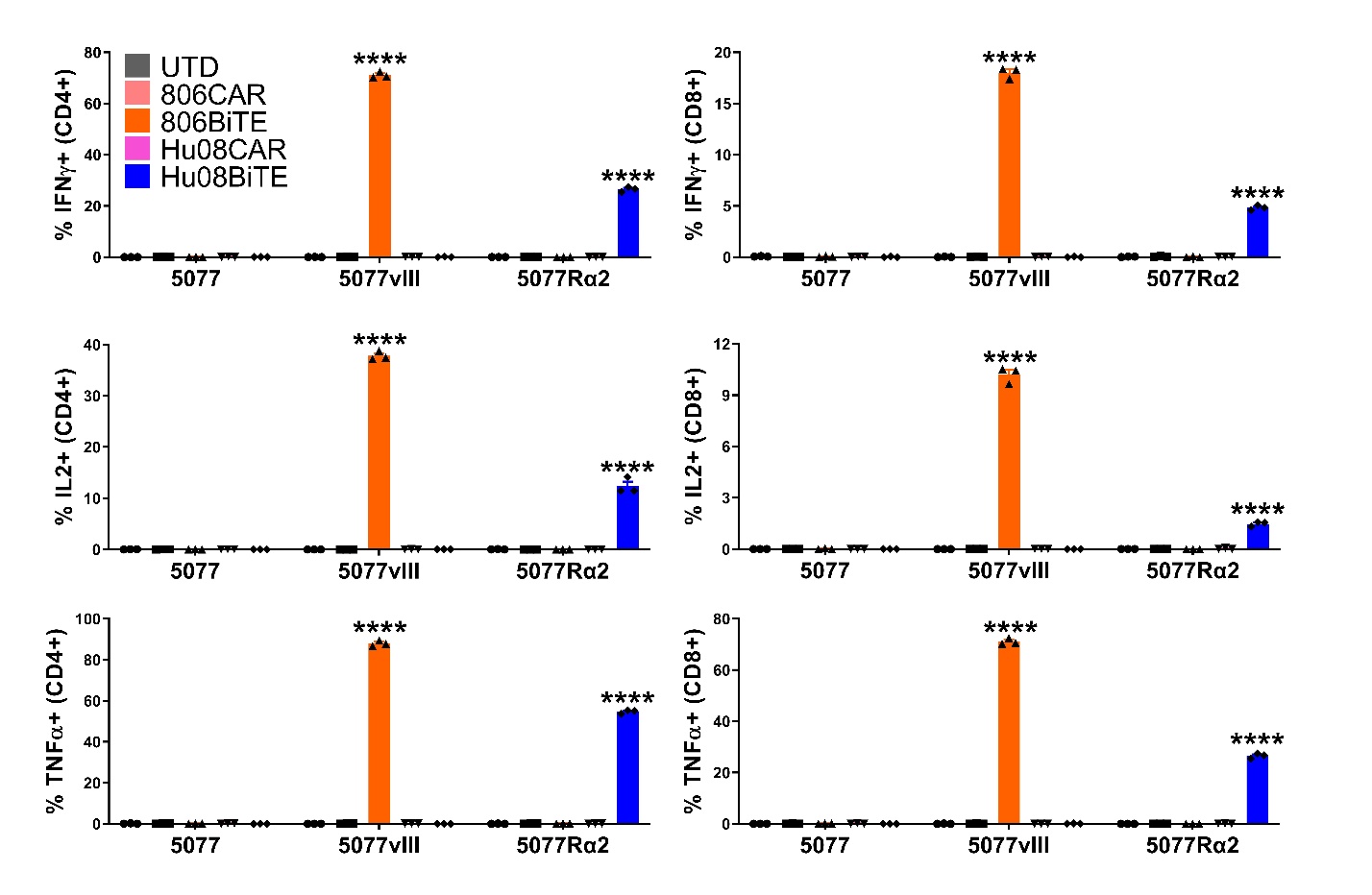

**Figure S1. BiTEs significantly responding to target positive glioma cells**

Flow based intracellular cytokine (IFNγ, IL2 and TNFα) staining of UTD T cells (UTD) co-cultured with target cells in conditioned media of CAR/BiTE T cells. CD4+ (left) and CD8+ (right) subgroups of T cells were distinguished by human CD8 staining. Statistically significant differences were calculated by one-way ANOVA with post hoc Tukey test. ****p<0.0001. Data are presented as means ± SEM.
